# Classic machine learning on top of multiple position weight matrices improves genomic prediction of transcription factor binding sites

**DOI:** 10.64898/2026.05.12.724515

**Authors:** Pavel Kravchenko, Ilya E. Vorontsov, Vsevolod J. Makeev, Ivan V. Kulakovskiy, Dmitry D. Penzar

**Author notes:** **Supplementary information** Supplementary data are available at github (https://github.com/autosome-ru/ArChIPelago) and at Zenodo (doi:10.5281/zenodo.14927304).

## Abstract

**Motivation:** DNA motifs recognised by transcription factors are typically represented as position weight matrices (PWMs), assuming independent contributions of individual nucleotides to protein binding specificity. Many alternative models accounting for correlations of positional contributions have been introduced in the past decades. However, performance gains have generally not outweighed the advantages of simplicity, interpretability, and practical applicability of PWMs with the well-established codebase. Existing software tools and motif databases provide multiple non-identical PWMs for the same transcription factor or even for the same dataset. It remains a practical question whether these PWMs can be effectively combined into a single improved model.

**Results:** Here we describe ArChIPelago (https://github.com/autosome-ru/ArChIPelago), a computational framework that combines multiple PWMs into a joint model using classic machine learning techniques, from linear regression to ensembles of decision trees. We show that such a combination improves prediction of transcription factor binding sites in genomic sequences. With a diverse collection of 704 ChIP-Seq datasets spanning 36 orthologous human and mouse transcription factors of diverse structural families, we show that ArChIPelago consistently outperforms the best available individual mono- and dinucleotide PWMs as well as sparse local inhomogeneous mixture models. Furthermore, using both human and mouse data, we demonstrate that PWM ensembles are capable of making reliable cross-species predictions.

## 1 Introduction

The morphology and function of a living organism are largely determined by genes, whose expression is coordinated by gene regulatory networks (Guo 2014), starting with the transcriptional level. Many biological processes are regulated in this manner, including embryonic development (Harrison et al. 2011), cell differentiation (Liu et al. 2008), and immune response (Roy 2019). Essential contributors to transcription regulation are transcription factors (TFs), which comprise a special class of DNA-binding proteins that can specifically bind to genomic loci (Kuiper et al. 2022) carrying occurrences of the respective DNA motifs, and affect transcription of functionally related genes.

Experimental and computational identification of TF binding sites (TFBS) has remained a challenge for several decades (Levy and Hannenhalli 2002; Zhu et al. 2002; Jayaram et al. 2016; Ladunga 2010), with significant progress being achieved with the adoption of high-throughput omics methods and bioinformatics techniques (Tognon et al. 2023; Dutta et al. 2026).

The classic way to represent the DNA motif is the position weight matrix (PWM, also known as position-specific scoring matrix, PSSM) (Stormo et al. 1982) (Stormo 2000; Schneider et al. 1986; Hertz and Stormo 1999). Since its adoption by Stormo, Berg, and von Hippel more than 30 years ago (Stormo et al. 1982; Berg and von Hippel 1987), it remains one of the most popular approaches not only for modelling TF-recognised DNA patterns (constructed via de novo motif discovery) but also for computational screening (motif search) for TFBS in genomic or synthetic sequences of interest (Ambrosini et al. 2020). The PWM is constructed under the assumption of independent additive contributions of neighbouring nucleotides to the total binding energy. Consequently, each position of a binding site can be unambiguously mapped to the respective column of a gapless multiple local alignment of known TFBS. This simplification enables construction of a convenient yet powerful model that provides sufficient reliability in TFBS prediction in numerous applications. To account for non-linear contributions of neighbouring or distant DNA bases to the binding specificity or to accommodate possible motif subtypes many alternatives to PWMs were proposed: dinucleotide PWMs (diPWMs) (Kulakovskiy et al. 2013; Segal et al. 2006), hidden Markov models (HMMs) (Raman and Overton 1994), sparse local inhomogeneous mixture (Slim) models (Grau et al. 2013), k-mer based support vector machines (Ghandi et al. 2014), and deep neural networks (Alipanahi et al. 2015; Bao et al. 2019). However, standard mononucleotide PWMs (monoPWMs) continue to be widely used in practice because of their simplicity, interpretability, and the ease of use with diverse existing software for PWM derivation and sequence scanning (Vorontsov et al. 2025; Boytsov et al. 2022). Further, they still provide a sufficiently hard-to-beat baseline for advanced models in joint benchmarking (Gryzunov et al. 2025).

The differences in experimentally assessed TF binding (whether from different experimental assays or conditions of the experiments) inevitably lead to variations in motif models produced in different studies or for different cell types. The experiment-to-experiment variations often arise from methodological and algorithmic limitations. Yet, various TFs can have intrinsically different binding modes resulting in distinct motif subtypes (Bais et al. 2011; Reid et al. 2010). Thus, for some TFs, the pattern of binding specificity cannot be precisely described by a single PWM. For instance, prediction of TP53 binding sites in model enhancers demonstrated a better performance for combinations of multiple PWMs (Verfaillie et al. 2016). Yet, it remains an open question whether combining multiple PWMs can be universally applicable for different TF families and motifs and whether the resulting complex models would learn the TF-specific but not experiment-, cell type-, or species-specific features of TF binding specificity.

Various benchmarking layouts have been proposed (Gryzunov et al. 2025; Vorontsov et al. 2025; Ambrosini et al. 2020) to select the single best-performing representation of the TF motif from the pool of PWMs produced by alternative motif discovery tools. However, it is tempting to utilise multiple alternative PWMs simultaneously, especially knowing that existing motif discovery tools yield similar but non-identical PWMs even from the same experimental dataset. Likewise, databases such as JASPAR (Rauluseviciute et al. 2024), HOCOMOCO (Vorontsov, Eliseeva, et al. 2024), and CIS-BP (Weirauch et al. 2014) in general contain many similar but non-identical PWMs for the same TF. An early attempt to combine PWMs with decision trees (Bi et al. 2011) using the model initialized with a single seed PWM with subsequent incorporation of ChIP-Seq (chromatin immunoprecipitation followed by deep sequencing) experimental data was partly successful, but showed unstable results and, in some cases, failed to outperform individual PWMs.

From the biological point of view, selection of the best-performing PWM could lead to exclusion of relevant binding modes. Owing to the rapid development of experimental in vivo and in vitro (Vorontsov, Kozin, et al. 2024) methods and cheaper high-throughput sequencing, thousands of large-scale genome-wide TF binding profiles from ChIP-Seq became available in the public domain, from the rigorous approach of ENCODE (The ENCODE Project Consortium 2012) to large-scale reanalysis of scattered public data in GTRD (Kolmykov et al. 2021), ReMap (Chèneby et al. 2020), and ChIP-Atlas (Hammal et al. 2022; Zou et al. 2024). The scale and diversity of these data enable training and evaluation of complex yet interpretable, lightweight, and easy-to-use models for TFBS classification and genomic TFBS prediction using classic machine learning techniques. Further, it becomes possible to evaluate if the prediction quality improvement remains with the model trained and tested on orthologous TFs of different species.

In our recent publication (Vorontsov et al. 2025), we showcased ArChIPelago, a model integrating predictions of multiple PWMs into a single machine learning model that demonstrated improved predictions for ChIP-Seq and genomic high-throughput sequencing systematic evolution of ligands by exponential enrichment (HT-SELEX) data. Here, we further explore this direction by utilising 704 ChIP-Seq datasets covering 36 orthologous human-mouse TF pairs with corresponding PWMs available in HOCOMOCO v11 (Vorontsov, Eliseeva, et al. 2024) to train binary classifiers using Logistic Regression, Random Forest, and Gradient Boosting. We show that aggregation of classical mono- and advanced dinucleotide PWMs leads to enhanced classification in comparison to the best existing individual PWMs, Slim models (Grau et al. 2013), or PWMs generated de novo from the entire training data. Interestingly, the inclusion of Slim models (Grau et al. 2013) of different orders as features yielded an additional improvement in prediction quality, surpassing that of single Slim models. Overall, we propose a simple and robust solution that improves the quality of PWM-based TFBS prediction, efficiently utilises multiple existing alternative PWMs as individual models combined in a unified classifier, overcoming the shortcomings of existing approaches that select a single PWM or account for a particular motif subtype.

## 2 Methods

### 2.1 Data preparation

ChIP-Seq datasets for *Homo sapiens* and *Mus musculus* transcription factors were obtained from GTRD (Yevshin et al. 2017) (Gene Transcription Regulation Database). The ChIP-Seq data were further filtered to consider only TFs with at least 3 datasets for each species and only experiments with matched control data, resulting in 414 human and 290 mouse ChIP-Seq datasets (Sup. Table 1). The peak calls made with MACS (Zhang et al. 2008) using hg38 *Homo sapiens* and mm10 *Mus musculus* reference genomes had been preprocessed during preparation of the HOCOMOCO v11 motif collection (Vorontsov, Eliseeva, et al. 2024), where the peak lists were filtered by “`tags” >= 10` and sorted by - `log10(P value)`. Additionally, the repeat-overlapping peak calls were removed with pybedtools (Dale et al. 2011) `subtract (f=0.7, N=True)` using RepeatMasker (Smit, AFA, Hubley, R & Green, P. 2013-2015) annotation. For each peak, the putative binding region was extracted as 301bp [-150;+150] centred at the peak summit.

PWMs used in this study were obtained by motif discovery from the same sets of GTRD peaks (a median of 14099 peaks per TF for human and mouse together) and computationally and manually curated according to the procedure used to obtain genuine motifs of the respective TFs for HOCOMOCO v11 motif collection (Vorontsov, Eliseeva, et al. 2024). As a result, we selected 36 TFs for which there were at least three classic mononucleotide position weight matrices (monoPWMs) and 3 dinucleotide PWMs (diPWMs) both for *Homo sapiens* (a median of 17 PWMs per TF) and *Mus musculus* (a median of 16–17 PWMs per TF) (Sup. Table 2). The complete preprocessed data and PWMs are available at Zenodo [10.5281/zenodo.14927304].

### 2.2 Data splitting and feature generation

The positive class label (1) was assigned to sequences encompassing the ChIP-Seq peak summits of the TF. To minimise computation time, up to 10,000 positives per TF were randomly sampled without replacement from all available filtered peak subsets (Sup. Table 2). Up to 1,000,000 negatives (yielding a class balance of 1:100) were selected for each TF from the list of peaks of all other TFs from unrelated TF families according to TFClass (Wingender et al. 2018). Additionally, we controlled the GC composition of the negative sequence set to match that of the positives using BiasAway (Khan et al. 2021). The ArChIPelago features were obtained by scanning genomic regions with each PWM using SPRY-SARUS (Kulakovskiy et al. 2016): the log-odds best hits of each PWM in each sequence were identified using `--skipn --show-non-matching --output-scoring-mode score besthit`.

### 2.3 Feature preprocessing and model performance assessment

Features were composed into a feature matrix with class labels (1, 0). The matrix was scale-transformed with sklearn.preprocessing.StandardScaler (scikit-learn 1.3.2 (Pedregosa et al. 2011)) and split into the training and testing sets independently for *Homo sapiens* and for *Mus musculus* in approximately a 30:70 test-to-train ratio. Following the design from (Zhou and Troyanskaya 2015), chromosomes 1, 8, and 21 were reserved for testing, and the remaining chromosomes (excluding 1, 8, and 21) were used for training for *Homo sapiens*. A similar set was built for *Mus musculus* with chromosomes 1, 8, and 19 reserved for testing. Sex chromosomes were excluded from the analysis. To estimate the ArChIPelago performance, we computed the area under the receiver operating characteristic (auROC) and the area under the precision-recall curve (auPRC) with the PRROC R package (Grau et al. 2015).

### 2.4 Model parameters

Using training data, we utilised the basic sklearn GridSearchCV module for parameters fitting resulting in the best performance of RandomForestClassifier (ArChIPelago) with `max_depth: 6, max_samples: 0.8, n_estimators: 100`; XGBClassifier with `colsample_bytree: 0.8, gamma: 0.3, max_depth: 3,min_child_weight: 1, n_estimators: 100,reg_alpha: 0.01, subsample: 0.8;`BaggingClassifier with XGBClassifier as a base classifier `colsample_bytree: 0.8, gamma: 0.3, max_depth: 3, min_child_weight: 1, n_estimators: 100, reg_alpha: 0.01, subsample: 0.8;` BaggingClassifier with LogisticRegression as a base classifier `C: 0.1, penalty: l2, solver: liblinear`; BaggingClassifier with RandomForestClassifier as a base classifier `max_depth: 25, max_samples: 0.8, n_estimators: 300.`

### 2.5 Slim models

The application of sparse local inhomogeneous mixture (Slim) models follows the logic of (Grau et al. 2013). In brief, the models were trained and tested side-by-side with ArChIPelago. Extended genomic regions of 1001 bp around the same ChIP-Seq peak summits as used for ArChIPelago were extracted and used to train the Slim models, including the peak and signal annotations required for Slim training, where signal was derived from the -10*log10(pvalue) field of the MACS output. The following parameters were used for training: `markov_order = 0` (equivalent to monoPWM), or `1` (equivalent to diPWM), or `-5` (to construct the so-called *limited* sparse local inhomogeneous mixture (LSlim) models (Keilwagen and Grau 2015) with a maximum distance of considered dependencies equal to the specified negative value), `background_markov_order = -1`, and `weighting_factor = 0.2`. Slim models were trained with background information comprised of the same set of negative examples used for ArChIPelago training. `Max score` was used to compute performance. The implementation of the Slim Python wrapper is available at GitHub (https://github.com/autosome-ru/ArChIPelago).

### 2.6 diChIPMunk models

DiPWMs from the full set of “positive” sequences (as used for training the ArChIPelago models) were constructed for each of the TFs with diChIPMunk: `ruby run_dichiphorde8.rb{output_directory} 15:15 filter y 1.0 m:{train_file.fasta} 200 20 1 4 random auto flat` (Kulakovskiy et al. 2010).

### 2.7 Data and code availability

The processed ChIP-Seq datasets and PWMs are available in the ArChIPelago repository on GitHub. The ArChIPelago source code and generated models are available on GitHub (https://github.com/autosome-ru/ArChIPelago). Peak coordinates and sequences for the training and testing data sets, PWMs, ArChIPelago models, and metadata are available on Zenodo (doi:10.5281/zenodo.14927304).

## 3 Results and discussion

### 3.1 Overview of the ArChIPelago workflow

ArChIPelago (https://github.com/autosome-ru/ArChIPelago), an ensemble framework combining multiple PWMs with ChIP-Seq data, employs machine learning for improved prediction of transcription factor binding sites (Fig. 1). We utilise ensemble learning to capture multiple binding motif subtypes and improve TFBS prediction performance across diverse TF families. We trained the ArChIPelago model on human data and tested it on human and mouse data, demonstrating cross-species transferability. Overall, for a wide range of TFs assayed with ChIP-Seq, ArChIPelago demonstrates improvements in the prediction quality over established TFBS prediction methods.

**Figure 1.**
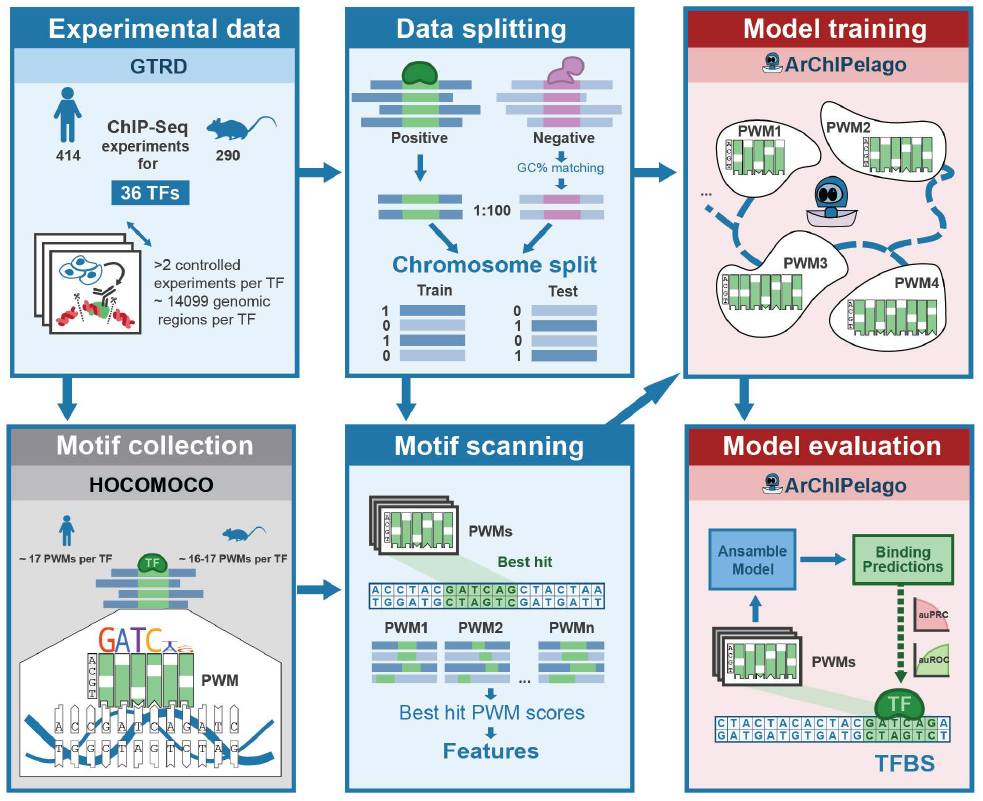
An overview of data preprocessing, ArChIPelago model design, training, and evaluation. ChIP-Seq data from the GTRD database for 36 TFs with more than 2 experiments in both mouse and human are selected. These data are used to create train-test splits and perform motif scanning using multiple PWMs per TF. Finally, the ArChIPelago ensemble model is trained on the PWM’s best hit scores and tested on a held-out dataset.

Utilising the evolutionary conservation of DNA-binding domains and respective DNA motifs, we assembled a rich set of 704 ChIP-Seq datasets encompassing 36 human-mouse pairs of TFs from diverse structural classes using the intermediate data from HOCOMOCO v11 (Vorontsov, Eliseeva, et al. 2024), enabling cross-species validation. We also collected 1558 mono- and di-nucleotide human PWMs (with a median of 17 PWMs per TF) and 1253 mouse PWMs (with a median of 16-17 PWMs per TF). Genomic features for subsequent model training were obtained by scoring sequences with the collected PWMs. ArChIPelago models used non-overlapping GC-matched peak sets with particular chromosomes dedicated exclusively for training or testing (see Methods).

### 3.2 ArChIPelago outperforms individual mono- and di-nucleotide PWMs

By design, ArChIPelago integrates multiple PWMs predictions into a joint TF-specific classifier of TFBS (positives, bound by the TF) and non-bound (negatives) sequences. Our ensemble approach surpasses the single best PWM by integrating predictions from similar yet complementary models, which may represent, e.g., alternative binding motif subtypes (Vorontsov, Eliseeva, et al. 2024; Lambert et al. 2018). First, we focused on the human ChIP-Seq data.

Within the ArChIPelago framework, we tested multiple models: a Random Forest (RF), Gradient Boosting (XGBoost), Logistic Regression, and Bagging (with Gradient Boosting and Logistic Regression as the base models). In all cases, ArChIPelago models systematically outperformed individual best available PWMs (Fig. 2), which were selected from the available pool by maximising auROC and auPRC on the train set.

**Figure 2.**
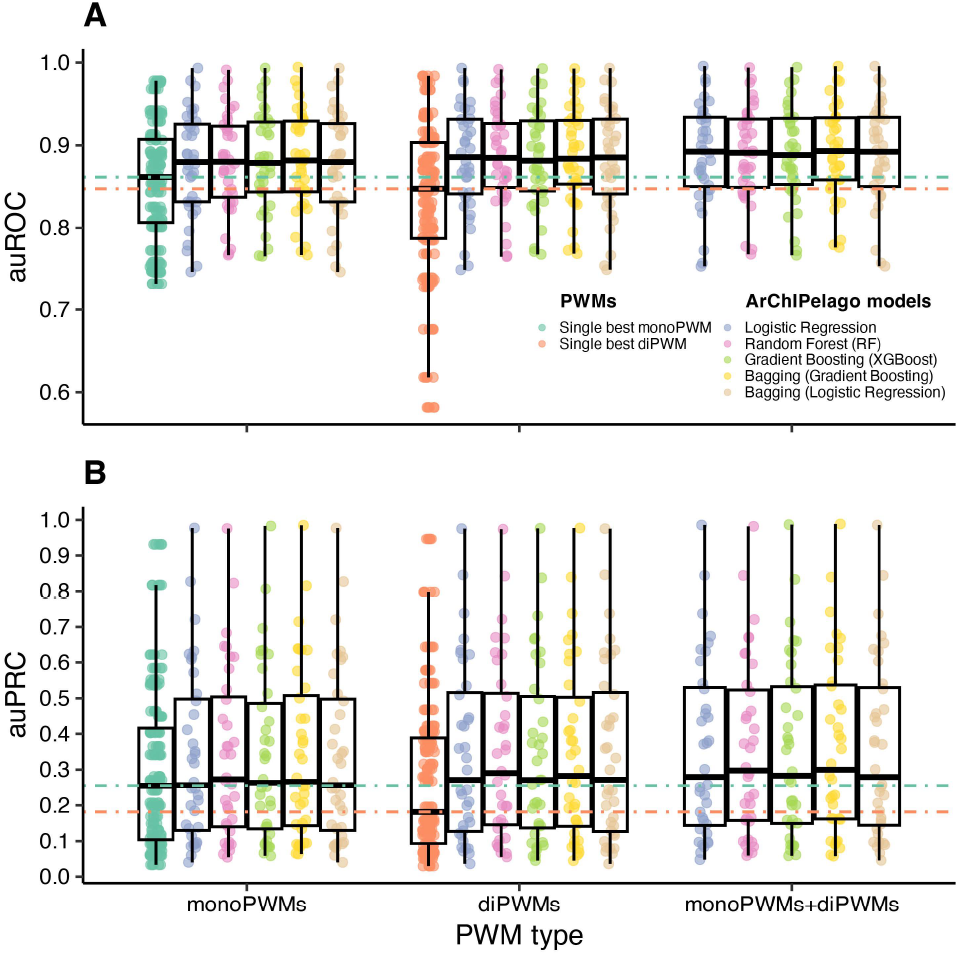
Integrative ArChIPelago models constructed from PWMs predictions outperform individual PWMs. Random Forest (RF), Gradient Boosting (XGBoost), Logistic Regression, and ensemble Bagging, trained on features composed from PWM predictions, consistently achieve superior performance compared to both best monoPWMs (green horizontal line: average across TFs) and best di-PWMs (orange horizontal line: average across TFs). Each dot represents a model performance for a single TF (A is auROC, B is auPRC). For a fair comparison, the best PWMs are selected on the train set, independently for comparing auROC and auPRC. The single best monoPWM and single best diPWM performances are shown only once in the corresponding categories for comparison.

Notably, despite consistent improvements over individual PWMs and high auROC, the absolute auPRC values remain moderate for most transcription factors (Fig. 2). This behavior is expected and reflects the intrinsic difficulty of TFBS prediction from genomic sequence alone under a strong class imbalance (1:100). In this context, modest gains in auPRC represent meaningful improvements in ranking true binding sites among a vast excess of negatives. Well-characterized transcription factors with long and information-rich motifs, such as CTCF, constitute an illustrative exception: for these TFs, even a single high-quality PWM already achieves high predictive performance, leaving limited room for improvement through model aggregation (Sup. Table 3). In contrast, for the majority of TFs with shorter, degenerate, or multi-modal binding motifs, ArChIPelago provides substantial and consistent gains, highlighting the benefit of integrating multiple complementary motif representations.

The consistent performance improvement across RF, Gradient Boosting, and Logistic Regression demonstrates that the benefits arise from integration of multiple basic models rather than algorithm-specific optimisation advantages. This is further illustrated by the simultaneous improvements in both auROC (Fig. 2A) and auPRC (Fig. 2B), indicating that the ArChIPelago genuinely enhances TFBS detection accuracy across multiple TF families rather than technically optimises a particular performance measure. While all models show comparable auROC improvements, RF achieves the slightly higher auPRC values, making it the method of choice for consequent analysis. In summary, multiple classification algorithms validate the advantage of the ArChIPelago ensemble principle and highlight the choice of the RF classifier as a base “to go” model.

If we limit the model to classic PWMs (monoPWMs) or diPWMs, the results are fully consistent across different classification models. Interestingly, individual diPWMs yield inferior predictions compared to mono-PWMs (Fig. 2). However, despite lower individual performance, diPWMs as a part of ArChIPelago models contributed significantly to the prediction quality. This enhancement suggests that dinucleotide dependencies capture critical aspects of protein-DNA recognition that complement mononucleotide preferences, but require an extra model layer to generalize to held-out data. Thus, to benefit from the higher basic prediction quality of monoPWM and the increased prediction quality of diPWM, we integrated both PWM types.

### 3.3 ArChIPelago outperforms individual PWM predictions by accounting for alternative motif subtypes

To study the ArChIPelago performance in more detail, we compared its predictions against the best monoPWM predictions on the level of individual TFs. According to both auROC and auPRC, ArChIPelago trained on only monoPWM, only diPWM, or on a mixture of monoPWMs and diPWMs predictions outperformed the best monoPWM for each of the tested TFs (Fig. 3A, B). The tight clustering of model performances above diagonal baseline references indicates that the ensemble approach benefits diverse transcription factor families rather than only specific binding architectures. ArChIPelago trained on monoPWMs and diPWMs demonstrated a median 0.022 increase in auROC compared to the best available monoPWMs (0.891 vs. 0.869, respectively) and a median 0.043 increase in auPRC (0.298 vs. 0.255, respectively).

**Figure 3.**
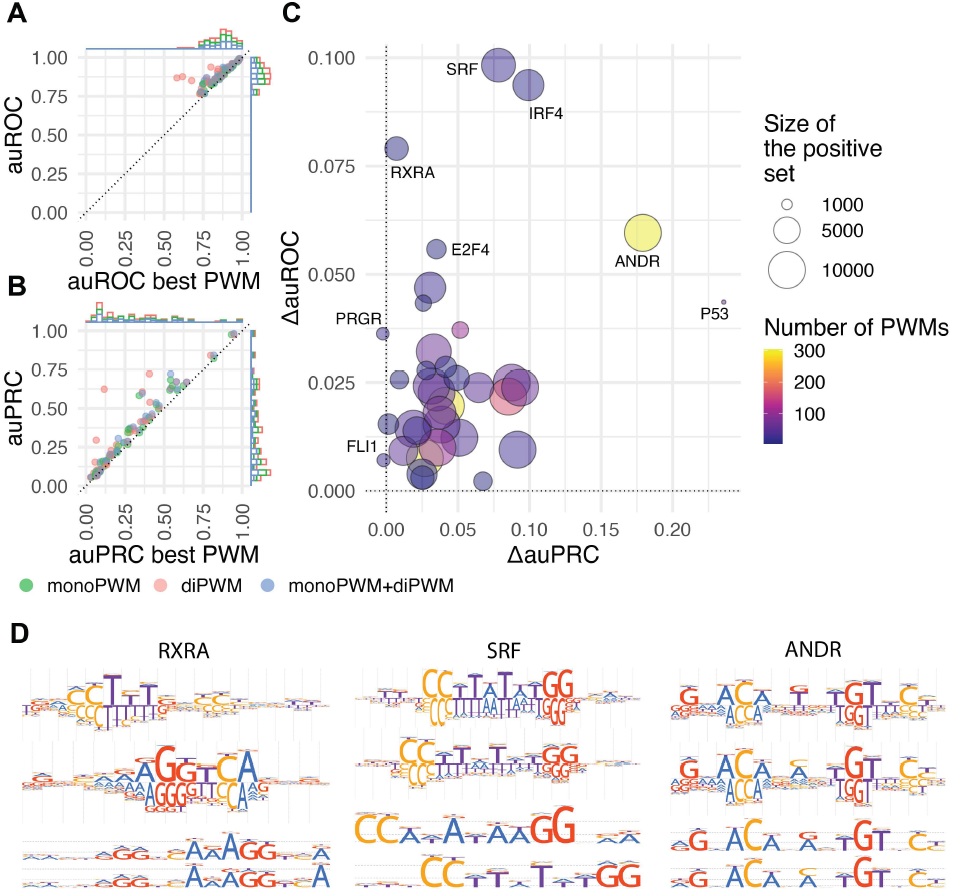
ArChIPelago outperforms the best PWMs by accounting for different motif subtypes. A. An overall auROC improvement for models built with several PWMs (mono, di, or both) in comparison to the best single monoPWM. Three colored dots per TF are shown. B. An overall improvement of auPRC: models built with several PWMs in comparison to the best single monoPWM. C. The difference in auROC and auPRC between the ArChIPelago model (trained on both monoPWMs and diPWMs) and the individual best monoPWM. The number of PWMs reflects the total number of monoPWMs and diPWMs for a given TF. D. Examples of PWMs reflecting alternative motif subtypes for RXRA, SRF, and ANDR (AR, Androgen receptor) TFs, which show the highest performance gain according to at least one of the metrics. The top 2 (ranked by the RF feature importance) monoPWMs and diPWMs are shown.

We then focused on the quantitative difference (ΔauROC, ΔauPRC) between RF models (built from monoPWMs and diPWMs) and the best monoPWM predictions for individual TFs (Fig. 3C). Overall, ArChIPelago models were able to outperform the best monoPWM. Only two TFs, FLI1 and PRGR (PGR, Progesterone receptor) showed no improvement as compared to the best PWM (technically, decreased auPRC) with ArChIPelago when trained and tested on human data, likely due to noise, unknown biases in the training data, or limitations of the PWM set. At the same time, 3 TFs SRF, IRF4, ANDR (AR, Androgen receptor) demonstrated greater than 0.05 increase in both auROC and auPRC. For some TFs (RXRA), the improvement was observed only for auROC. At the same time, there was an exceptional auPRC improvement (>0.2) with moderate auROC gains for P53, suggesting that ArChIPelago modelling captures complex context-dependent binding of this important TF (Vyas et al. 2017) across diverse genomic loci.

Although no clear global correlation was observed between the number of PWMs and performance gains, classifiers generally benefited from the inclusion of additional PWMs up to a TF-specific saturation point. The examples of RXRA, SRF, and ANDR demonstrate that the most dramatic increase in prediction quality was achieved for TFs with several distinct motif subtypes that a single PWM cannot capture (Fig. 3D), in agreement with our earlier observation reported in (Vorontsov et al. 2025). It highlights the need for more comprehensive binding specificity assays (such as in (Jolma et al. 2024)) and underscores the importance of diverse motif collections, such as HOCOMOCO (Vorontsov, Eliseeva, et al. 2024), CIS-BP (Weirauch 2014), and JASPAR (Rauluseviciute et al. 2024).

To ensure that ArChIPelago does not memorise but generalises the diversity of TFBS, we utilised a cross-species validation by training on human and testing on mouse ChIP-Seq data (Fig. S1, Methods). A similar but weaker performance gain of ArChIPelago models over the best PWMs to that observed in the within-species evaluation (Fig. 2) was also noted in the cross-species evaluation (Fig. S2). IRF4, SOX2, and ESR1 TFBS predictions were improved by more than 0.05 in auROC, and ANDR and ESR1 by more than 0.35 in auPRC (Fig. S3). At the same time, for 12 TFs, the ArChIPelago performance was worse than that of the best PWM according to at least one of the metrics, suggesting that the models relied on either species-specific or cell type-specific sequence features, as in our data, the set of ChIP-Seq cell types was not aligned between human and mouse. Overall, the best-performing TFs are partially shared between the within-species and cross-species evaluations, with the former demonstrating a higher performance gain.

## 4 ArChIPelago outperforms previous-generation TFBS models

To further explore the performance of ArChIPelago models, we benchmarked their predictions against the diPWM discovered by diChIPMunk (Levitsky et al. 2014) from the complete set of ‘positive’ examples, and Slim models (Grau et al. 2013) of different orders. Surprisingly, the individual Slim (m=0 equivalent to monoPWM, Slim m=1 equivalent to diPWM, LSlim m=-5) models built from complete positive genomic sequences show negative or near-zero delta auROC and auPRC values, indicating their inferior performance compared to traditional monoPWMs (first three lines in Fig. 4 A and B). Despite being trained on comprehensive positive sequence data, the diChIPMunk-derived diPWMs also fail to achieve superior performance over the best monoPWMs, highlighting the challenges of the past-generation approaches even with larger datasets.

**Figure 4.**
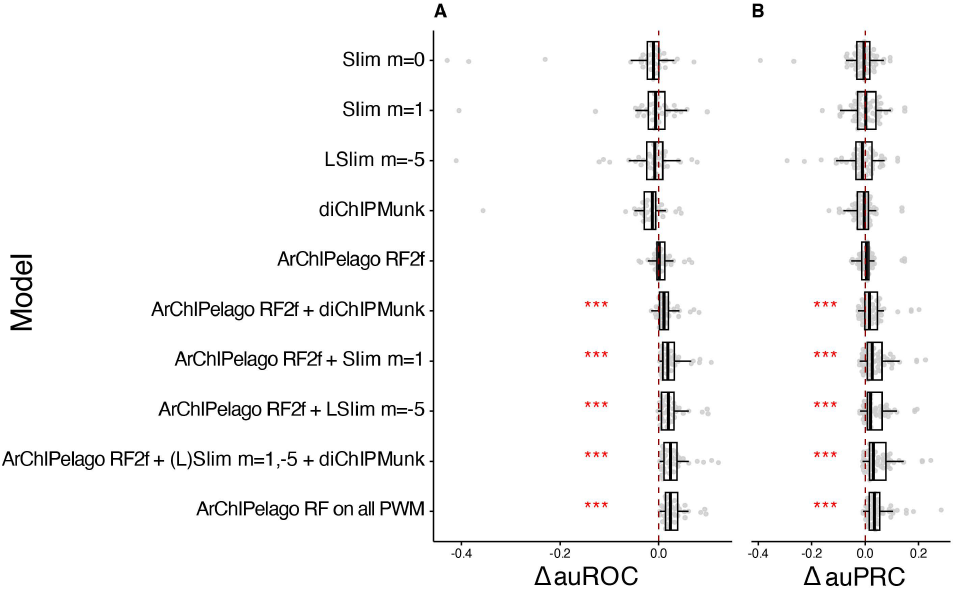
ArChIPelago outperforms diPWMs and Slim models of different degrees. Slim and diChIPMunk features improve ArChIPelago predictions. The comparison of ΔauROC (A) and ΔauPRC (B) between the best monoPWMs and individual models, namely Slim m=0 (equivalent to monoPWM), Slim m=1 (equivalent to diPWM), LSlim m=-5, diChIPMunk diPWM (built from all the positive genomic sequences), ArChIPelago RF on the best monoPWM and diPWM includes two features (RF2f, two features only): predictions of the best monoPWM and of the best diPWM, and combinations of the above-mentioned included into the RF as additional features. *** p < 0.001, Wilcoxon signed-rank test. Statistical significance was assessed for each variable group, comparing the distribution of values to a median of zero. Predictions of the human-trained model on the human test set are shown.

On the contrary, the predictions produced by the RF model built using only two matrices, the top-scoring monoPWM and diPWM (Fig. 4: RF on the best PWMs, two features only), as expected, were close to the individual best monoPWM predictions. Thus, with a minimal set of features, ArChIPelago replicates the baseline performance. The comparison of RF with the addition of Slim and/or diChIPMunk models (RF+Slim m=1 three features in total, RF+LSlim m=-5 three features in total, RF+diChIPMunk three features in total, RF+Slim m=1+LSlim m=-5+diChIPMunk five features in total) demonstrates the plasticity of the ArChIPelago framework and the progressive predictive advantage that more complex models give to the unified classifier. Notably, the performance of the RF trained on all available monoPWMs and diPWMs is comparable to or better than that of the TF models incorporating Slim and diChIPMunk features, suggesting that lightweight, publicly available PWMs may be sufficient for high-quality TFBS identification. A similar result was achieved in the interspecies experiment (Fig. S4). Additionally, the tight clustering of RF model performance improvement (ΔauPRC and ΔauROC) above zero demonstrates the reliability of the ensemble approach across diverse TF families, in contrast to the variable and predominantly negative deltas of particular individual models (Slim m=0, Slim m=1, LSlim m=-5, diChIPMunk). Overall, every RF model that integrates all available PWMs or available PWMs with advanced features shows the improvement compared to the single best PWM (p<0.05, Wilcoxon signed-rank test), leveraging the advantages of ensemble approaches over individual model optimisation and traditional PWMs.

## 5 Conclusions

In this study, we explored the ArChIPelago framework by summarising individual predictions of PWMs into a unified classifier. In contrast to earlier attempts to combine PWMs with decision trees (Bi et al. 2011), ArChIPelago models do not select a single PWM, but integrate TFBS predictions from all available PWMs as features. ArChIPelago demonstrates median improvements of 0.022 auROC and 0.043 auPRC compared to the single best PWMs. Moreover, the model is applicable in a cross-species way, as we demonstrated by using human-trained RF to predict TFBS in mouse ChIP-Seq data (Fig. S3, S4).

In this work, we further demonstrated plasticity and applicability of ArChIPelago to real-world cases, in line with previously understudied TFs with a broad range of diverse PWMs (Vorontsov, Kozin, et al. 2024).

ArChIPelago can be used for integrating multiple features and predictions (motif scans) of basic models, and readily available PWMs can facilitate achieving state-of-the-art performance through ensemble modelling, which, in turn, makes high-quality TFBS prediction accessible without requiring specialised motif discovery expertise or excessive computational resources. The demonstrated applicability across species and diverse TF families, combined with the availability of scanning tools (https://github.com/autosome-ru/ArChIPelago-TFBS-finder) and curated datasets, establishes ArChIPelago as a practically viable solution for broader use.

## Authors’ contributions

I.V.K., D.D.P., and V.J.M. designed and supervised the study. P.K. implemented and evaluated ArChIPelago. D.D.P. implemented the Slim wrapper for Python. P.K. drafted the manuscript, which was finalized with the feedback of all authors.

## Acknowledgements

We are deeply thankful to Ivo Grosse and Jan Grau, who assisted with training Slim models.

