## Supplementary figures 1-4 for "Classic machine learning on top of multiple position weight matrices improves genomic prediction of transcription factor binding sites"

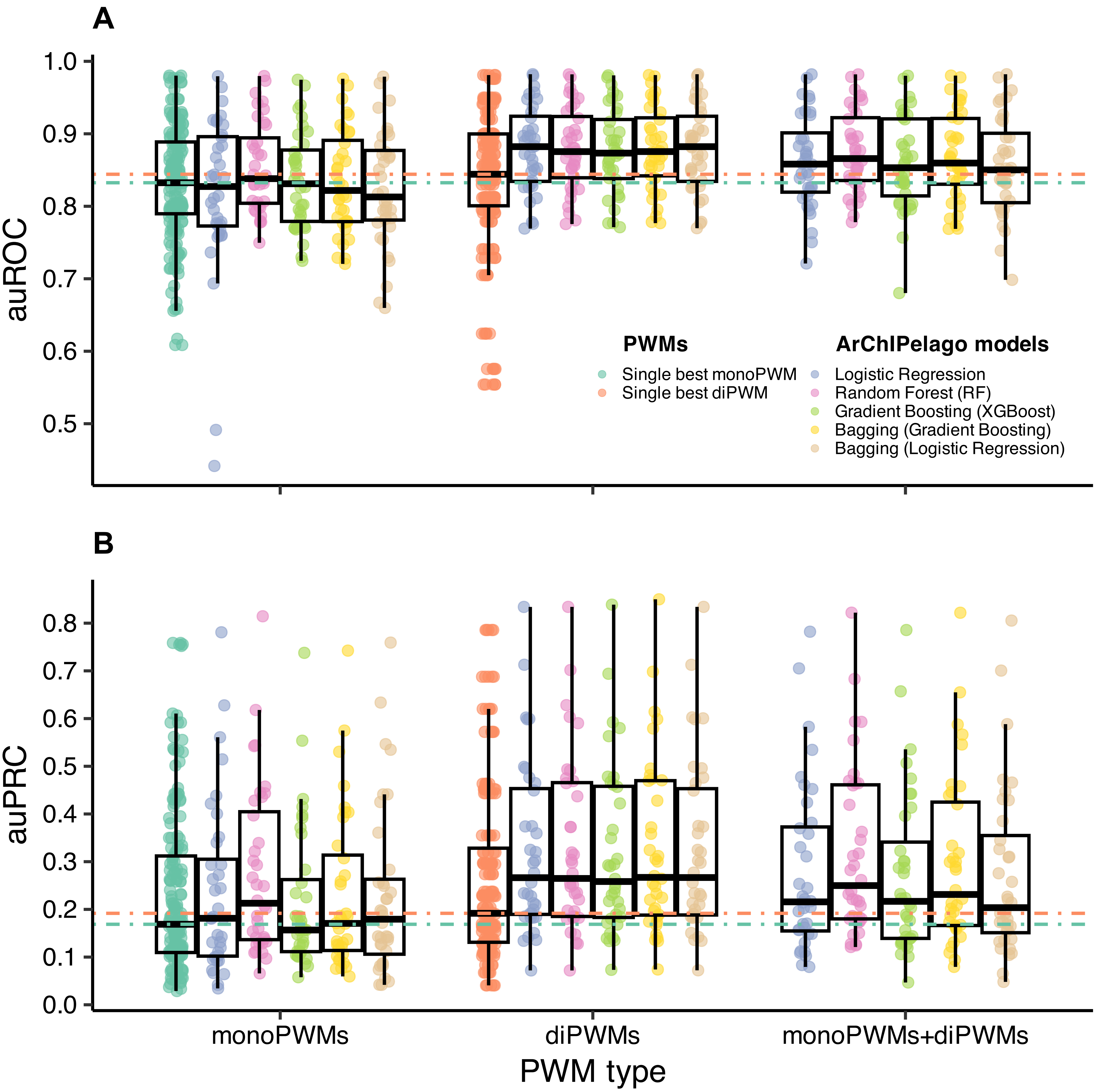


**Figure S1. Integrative ArChIPelago models outperform individual PWMs. The models were trained on human data and tested on mouse data.**
Random Forest, Gradient Boosting (XGBoost), Logistic Regression, and ensemble Bagging consistently achieve superior performance compared to both best monoPWMs (green horizontal line: average across TFs) and best di-PWMs (orange horizontal line: average across TFs). Each dot represents a model performance for a single TF (**A** is auROC, **B** is auPRC). For a fair comparison, the best PWMs are selected on the train set, independently for comparing auROC and auPRC.

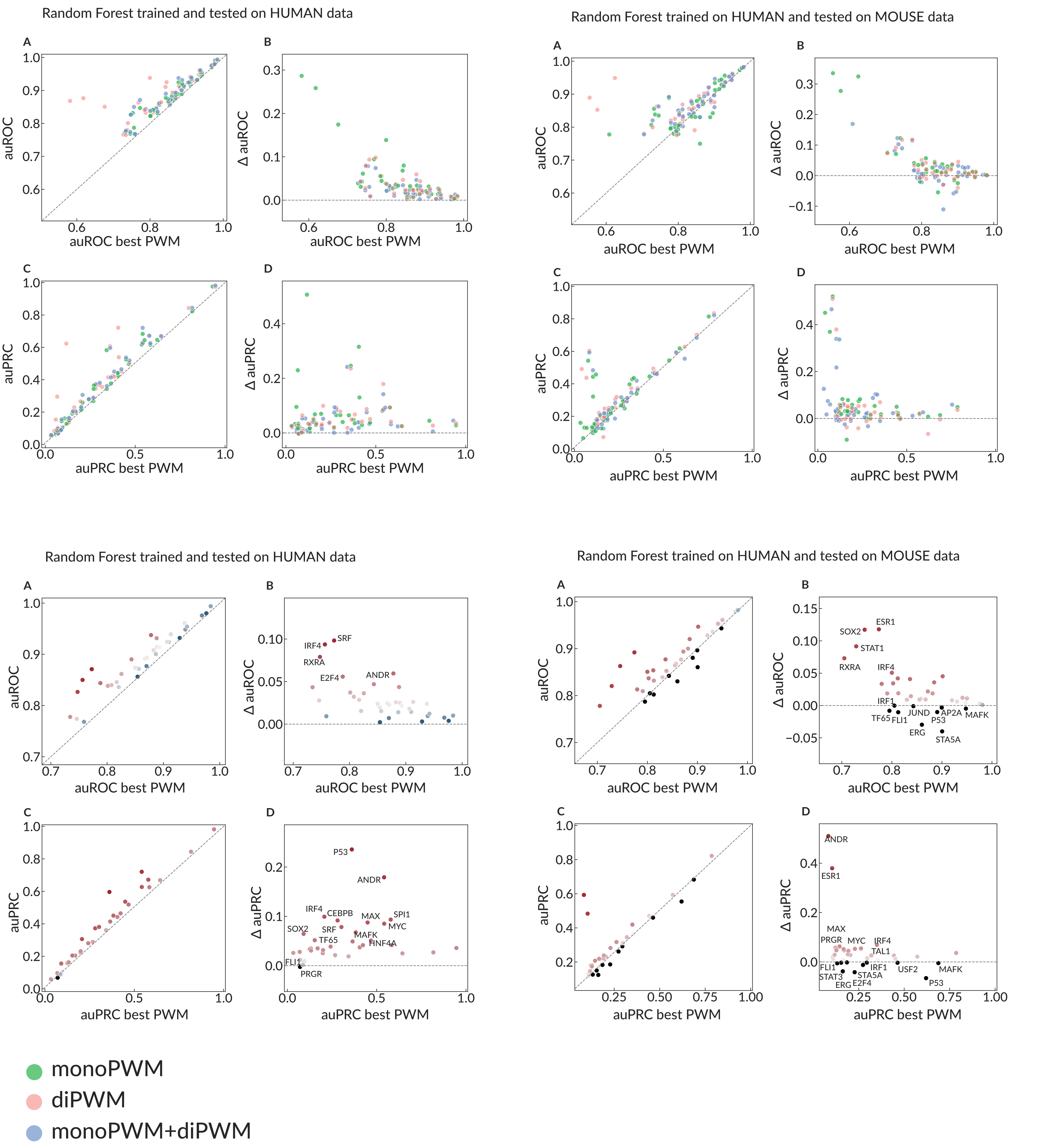


**Figure S2. Performance of the Random Forest trained and tested on human data (left column) and trained on human data but tested on mouse data (right column).**
**A, B** show auROC values and their differences (ΔauROC), while bottom panels **C, D** show auPRC values and their differences (ΔauPRC). Results are displayed for models trained with monoPWMs, diPWMs and combined monoPWMs and diPWMs (panels with all three PWM types combined) and for monoPWMs and diPWMs (panels with monoPWMs and diPWMs). Individual transcription factors are highlighted in the bottom row, illustrating cases with improved or reduced cross-species generalisation.

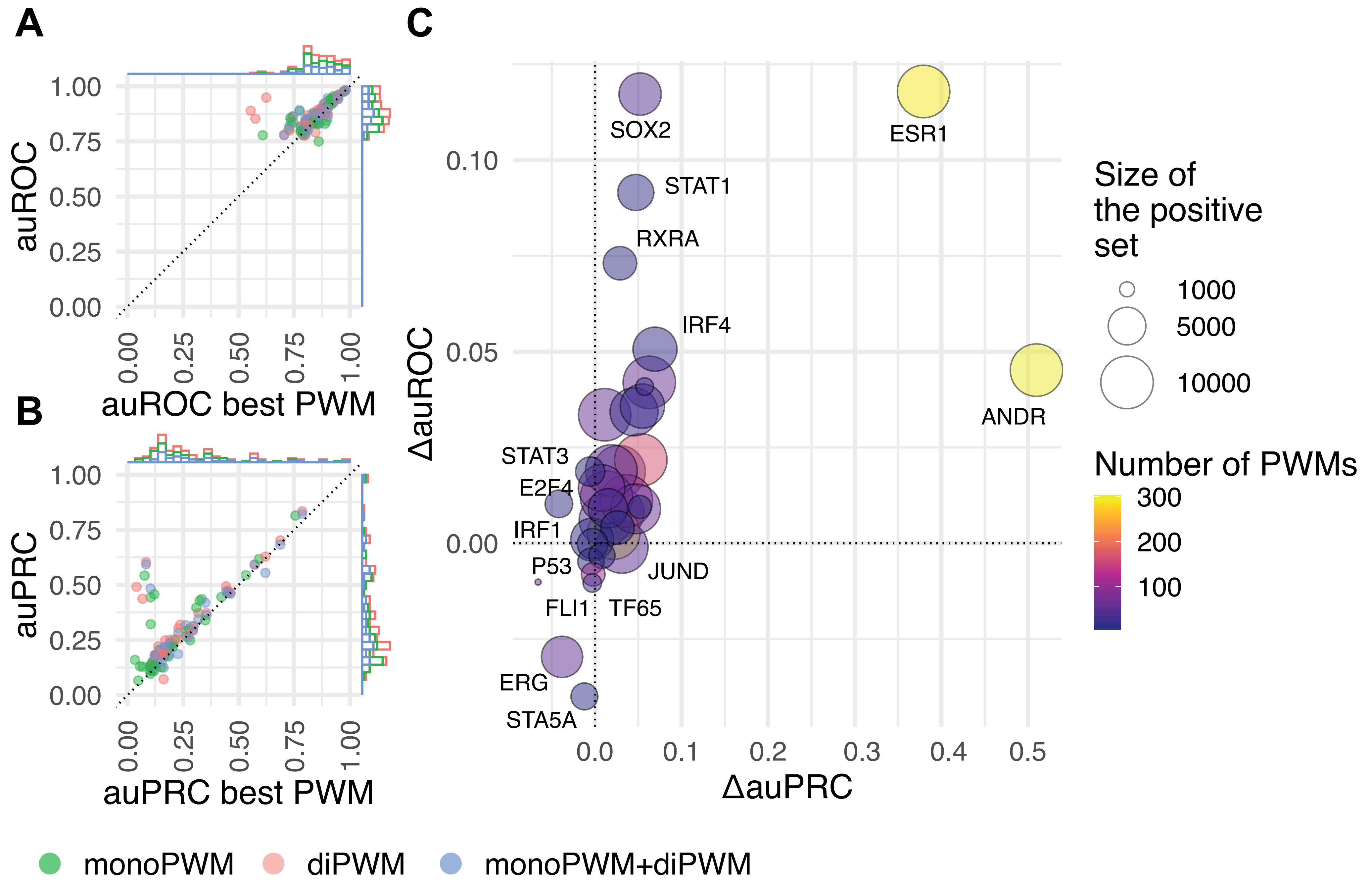


**Figure S3. ArChIPelago outperforms the best PWMs by accounting for different motif subtypes. The models were trained on human data and tested on mouse data.**

**A**. An overall auROC improvement for models built with several PWMs (monoPWMs, diPWMs, or both monoPWMs+diPWMs) in comparison to the best single monoPWM. Three colored dots per TF are shown. **B**. An overall improvement of auPRC: models built with several PWMs in comparison to the best single monoPWM. **C**. The difference in auROC and auPRC between the ArChIPelago model (trained on both monoPWMs and diPWMs) and the individual best monoPWM. The number of the PWMs reflects the total number of monoPWMs and diPWMs for a given TF.

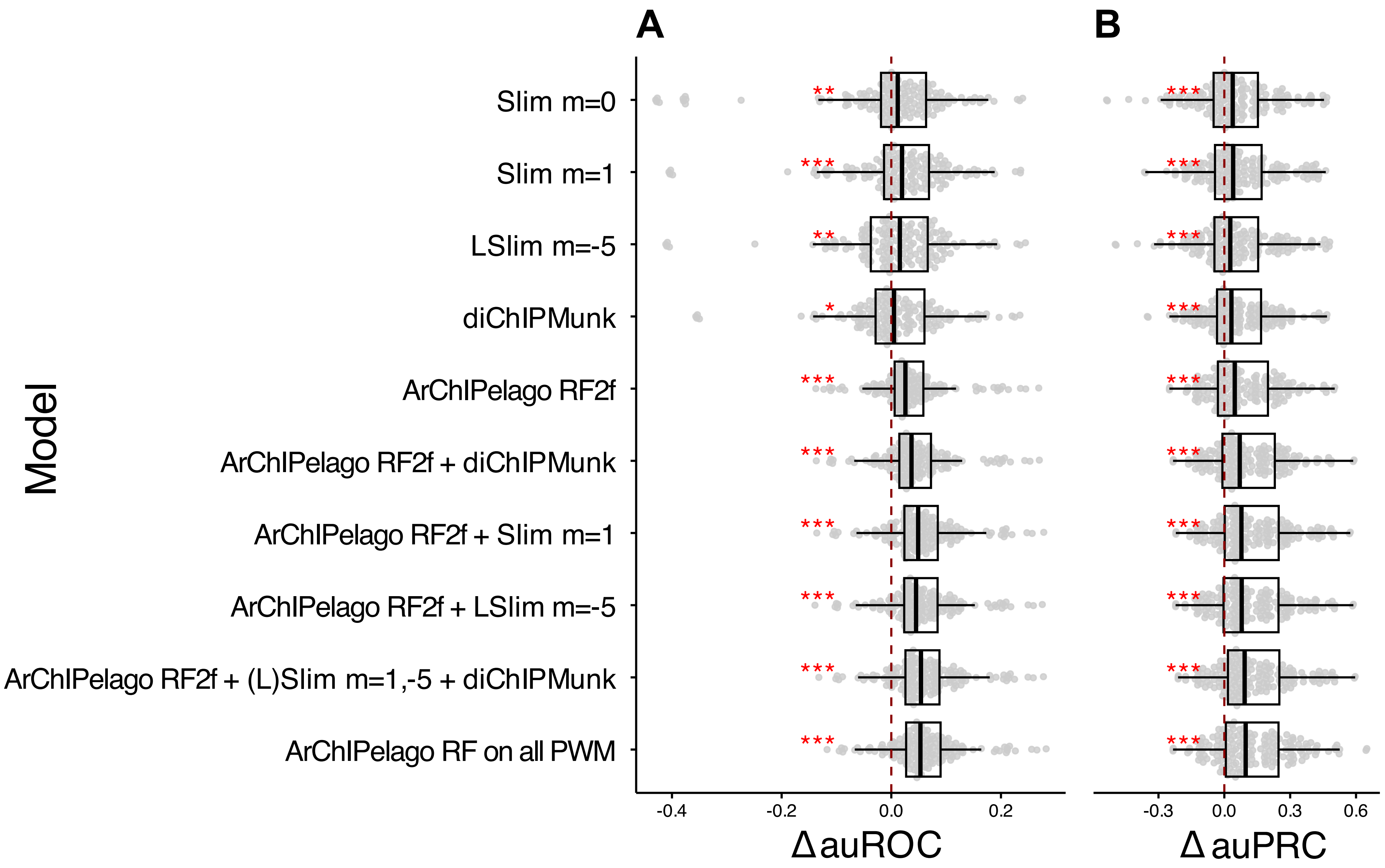


**Figure S4. ArChIPelago outperforms diPWMs and Slim models of different degrees. Slim and diChIPMunk features improve ArChIPelago predictions. The models were trained on human data and tested on mouse data.**

The comparison of ΔauROC (**A**) and ΔauPRC (**B**) between the best monoPWMs and individual models, namely Slim m=0, Slim m=1, LSlim m=-5, diChIPMunk diPWM (built from all the positive genomic sequences), ArChIPelago RF on the best monoPWM and diPWM includes two features: predictions of the best monoPWM and of the best diPWM (RF2f, two features only), and combinations of the above-mentioned included into the RF as additional features. Wilcoxon signed-rank test. Statistical significance was assessed for each variable group, comparing the distribution of values to a median of zero. One asterisk marks $p < 0.05$, two asterisks mark $p < 0.01$, three asterisks mark $p < 0.001$.

Supplementary tables

**Table 1.** The ChIP-Seq experiment identifiers selected for the model training. (https://github.com/autosome-ru/ArChIPelago/blob/main/Table_1.xlsx)

**Table 2.** The metadata for TFs selected for analysis: number of experiments, peaks, test and train sequences with the GC%. (https://github.com/autosome-ru/ArChIPelago/blob/main/Table_2.xlsx)

**Table 3.** The metadata for ArChIPelago performance. (https://github.com/autosome-ru/ArChIPelago/blob/main/Table_3.xlsx)
